# Neophilia in wolves and dogs

**DOI:** 10.1101/2025.04.24.650376

**Authors:** Dániel Rivas-Blanco, Lou Gonnet-dit-Revel, Friederike Range, Sabine Tebbich, Sarah Marshall-Pescini

**Author notes:** Both authors contributed equally to this work (Dániel Rivas-Blanco, Lou Gonnet-dit-Revel).

## Abstract

The study of domestication provides a unique opportunity to analyze the effects of natural selection in the attraction towards novelty —known as neophilia. This is chiefly due to the fact that selective pressures acting on domesticated animals are often greatly diminished or at least very different from their wild counterparts. In this study, we investigated the neophilic levels of three canine populations differing in their level of contact with human environments both from an evolutionary and ontogenetic perspective: wolves, pack-living dogs, and pet dogs. In order to study their neophilic response, we presented the animals with two objects. The first of these objects was displayed in the animals’ enclosure for several days. After the animals were habituated to this first object, the second one was introduced together with the first one in a shorter test session meant to explore their preferences for either of the objects. We predicted that dogs —and pet dogs in particular to display higher levels of neophilia, since human-created environments tend to change at a faster pace, and thus, a higher drive to explore these changes would be more beneficial than in a comparably more stable environment. Our results show no apparent differences between the populations in terms of latency to approach the new object, nor in the identity of the object they approached first. Nonetheless, all populations interacted more with the new object during the test phase. Wolves did also interact longer with the object presented in the first phase of the experiment.

## Introduction

Novel objects in the environment may offer new opportunities for an animal, but can also pose a potential risk. Neophobia and neophilia (respectively: fear of and attraction to novelty) are motivational factors” that greatly influence how animals interact with their environment (Griffin & Guez, 2014; Marshall-Pescini et al., 2017b). Neophilia and neophobia are not necessarily correlated with each other (Greenberg & Mettke-Hofmann, 2001), as the absence of a fear response towards a novel item does not necessarily imply an attraction toward it. Neophobia, can be detrimental to the potential exploitation of novel opportunities (Benson-Amram et al., 2012, Moretti et al., 2015), but can be advantageous (and thus selected for) in predator-rich or otherwise “dangerous” environments (Benson-Amram et al. 2012, Greggor et al., 2016b). Neophilia, on the other hand, allows an animal to easily gather information about changes in their environment (Tebbich et al., 2016). High levels of variability and complexity in an animal’s environment may select for neophilia, as interest towards a potentially new resource can have greater effects (Griffin & Guez, 2014; Mettke-Hofmann et al., 2002). Exploration persistence (i.e. the duration the animal spends investigating an object) is higher in species that need to process their food in order to remove inedible parts than in those that can access it directly (Mettke-Hofmann et al., 2002). Persistence is also often negatively impacted by neophobia (Biondi et al., 2010; Griffin & Guez, 2014; Rao et al., 2018) although in some species the opposite has been shown (see Moretti et al., 2015). In line with what would be predicted by the evolutionary pressures selecting for neophilia and neophobia, it has been found that a lower trophic position (hence a higher number of potential predators) begets higher levels of neophobia (Crane & Ferrari, 2017), while habitat variability and complexity correlates with higher levels of neophilia (Mettke-Hofmann et al., 2002).

Domestication provides an excellent model to study the evolution of both neophilic and neophobic traits. The effect of several natural selection pressures is diminished or overall different for domesticated animals. Most notable amongst these pressures is the absence of predation, which should lead to reduced neophobia. At the same time, human-created environments are more complex and prone to rapid change which should select for higher levels of neophilia (Griffin et al., 2016, Sih et al., 2012; Suzuki et al., 2021; but see Greggor et al., 2016a and Miranda et al., 2013).

Wolves and dogs may prove to be a particularly interesting model to test the effect of domestication on neophobia and neophilia due to their genetic closeness They only started to diverge around 30,000 years ago (Cooper et al., 2015), but occupy very different ecological niches. Wolves are cooperative hunters with a rather low success rate (between 10% and 50%; Freedman & Wayne, 2017; Mech et al., 2015). Conversely, dogs usually live in environments with an abundance of food either as pet dogs with humans directly providing food or as free-ranging animals, where the dogs scavenge on human refuse, with the latter scenario being likely the evolutionary context in which they thrived (Brooks, 1990; Butler et al., 2018; Butler & Du Toit, 2002). Considering this, the social-ecology hypothesis (Marshall-Pescini et al. 2017a) postulates that dogs should be more neophilic than wolves, as the anthropogenic niche would promote investigating new potential resources. Furthermore, wolves underwent severe persecution by humans (Dufresnes et al., 2018), thus there may have been a selection for shyer, more neophobic individuals who kept a distance from human settlements and human-created artifacts; (Range & Marshall-Pescini, 2022).

At the ontogenetic level, experience may influence how animals react towards novel objects. For example, captive hyenas are less neophobic and more explorative than their wild counterparts, implying that their experience with human-created constructs has had an effect on these motivational factors (Benson-Amram et al., 2013). Additionally, in dogs, it has been shown that persistence is higher in pet and pack-living captive dogs than in free-ranging dogs. It has been suggested that socialized dogs are encouraged by humans to interact with toys, which may have affected their overall object-directed motivation (Lazzaroni et al., 2019).

Traditionally, neophobia has been measured by placing food (or any other potential motivator) in a novel environment or within the vicinity of an unfamiliar item, and analyzing whether or not this motivator is approached (Greenberg & Mettke-Hofmann, 2001; Greggor et al., 2016a; Miranda et al., 2013). Neophilia, conversely, is often studied by the presentation of a novel object in a non-foraging context, with any potential approaches and interactions towards it being considered a measure of neophilia (as the animal would theoretically have no other reason to interact with it other than its novelty; Greenberg & Mettke-Hofmann, 2001; Heinrich, 1995; Miranda et al., 2013). Another way of testing neophilia is to investigate the preference an animal may have for a novel object over a known one (Kaulfuß & Mills, 2008). This approach has been used with pet dogs, with results showing that they preferentially interact with the novel object over the known one. The authors of the study suggested that neophilia may have been an adaptive response for ancient dogs, facilitating their approach to human settlements. However, without a direct comparison with wolves, this possibility remains untested.

Thus, the current study investigated the potential effects of domestication on neophilic behavior by comparing similarly raised and kept wolves and dogs. Furthermore, in order to test the effect of experience, pet dogs were also tested. Thus, we presented animals first with novel objects in their home enclosure (exposure phase) and, once habituated to the object, carried out a test in which, similarly to Kaulfuß and Mills, we simultaneously presented a new or the by-now-familiar object (test phase).

In a previous study by Moretti et al. (2015), dogs approached novel objects faster than wolves did, but wolves interacted with them for a longer amount of time. Based on these results and the socio-ecology of the two species, we expected pack dogs to approach and need less time to habituate to the object presented in the exposure phase than the wolves and to show a stronger preference towards the new vs. familiar object in the test phase. Conversely, we predicted that, while wolves would need more time to habituate to the new objects in both phases, they would also interact for longer with them than the dogs (see results from Moretti et al., 2015 and Rao et al., 2018), because they need to open and process the carcasses of their prey (Mech et al., 2015; Mettke-Hoffman et al., 2002). Additionally, pet dogs, being more familiar with human-made environments, were expected to interact more with the objects during the exposure phase than their pack-living counterparts (in line with Lazzaroni et al., 2019), and to show a preference for the new over familiar one (similarly to Kaulfuß & Mills, 2008).

## Methods

### Study area

We conducted the study at the Wolf Science Center (WSC), located in Ernstbrunn Wildpark in Lower Austria, from August 2021 to December 2021. We used 10 outdoor enclosures, ranging from 1000 m^2^ to 10000 m^2^ (see details table S1 in the Supplementary materials), all of them containing trees, bushes, and other naturally occurring items (stones, sticks, etc.).

To get a full view of the testing area, grass and branches were periodically removed around two selected trees within circles of a radius of 2.60 meters in wolf enclosures and 1.70 meters in dog enclosures. Additionally, we placed stones on the ground forming two concentric circles in order to calculate the latency of the animal’s approach towards the objects (more details in the ‘exposure phase’ sub-section).

### Subjects

In this experiment, we tested 6 pet dogs (1 female and 5 males), 6 pack-living dogs (4 females and 2 males; from 3 genetic lines) and 10 grey wolves (3 females and 7 males; from 6 genetic lines) at the WSC (which constitute all available subjects at the WSC at the time of testing; see information Table S2 in the Supplementary materials).

The pack dogs and the wolves were raised in a similar environment. They were all separated from their mothers at 10 days of age to be bottle-fed and hand-raised by humans for 5 months. During these months, the hand raisers did not initiate any sort of object-directed play with the pups, rather, the pups were left to play among themselves. Afterward, they were integrated into already existing packs or new packs were formed. The current packs are composed of two or three individuals that are not always related.

The pet dogs tested were the personal dogs of the trainers working at the WSC. The owners were informed about the main aim of the study and gave explicit approval to test their dogs in it and use the resulting data for scientific purposes. Despite the dogs belonging to different owners and living in different households, they had a good relationship with each other and were used to being kept together in an enclosure at the WSC without humans being present.

All subjects were familiar with being tested since they regularly participate in similar experiments. Furthermore, all three populations had similar ages (average age for wolves being 10.7 ± 0.77 years, 7.83 ± 0.54 years for pack dogs, and 8.50 ± 0.85 years for pet dogs).

### Test items

All the objects were built out of wood and painted with finger paint as it is non-toxic, and the animals were able to manipulate the objects with their mouths unrestrictedly. We used wood to build the objects because the wolves can easily break other materials, in a way that potentially endanger them. Moreover, when possible, we used blue and yellow paint as wolves and dogs only see a limited color spectrum, but their vision allows for differentiation of hues in the blue and yellow spectrums (Singletary 2021). We designed and built objects in pairs, matching the size, shape and color within the same pair as much as possible in order to reduce potential innate preferences towards any of these features (see Figure 1). We used one of the objects of each pair in the “exposure phase” of the experiment while both this object and the other object of the same pair were used for the “test phase” (see the “Experimental procedure” section for more details).

**Figure 1.**
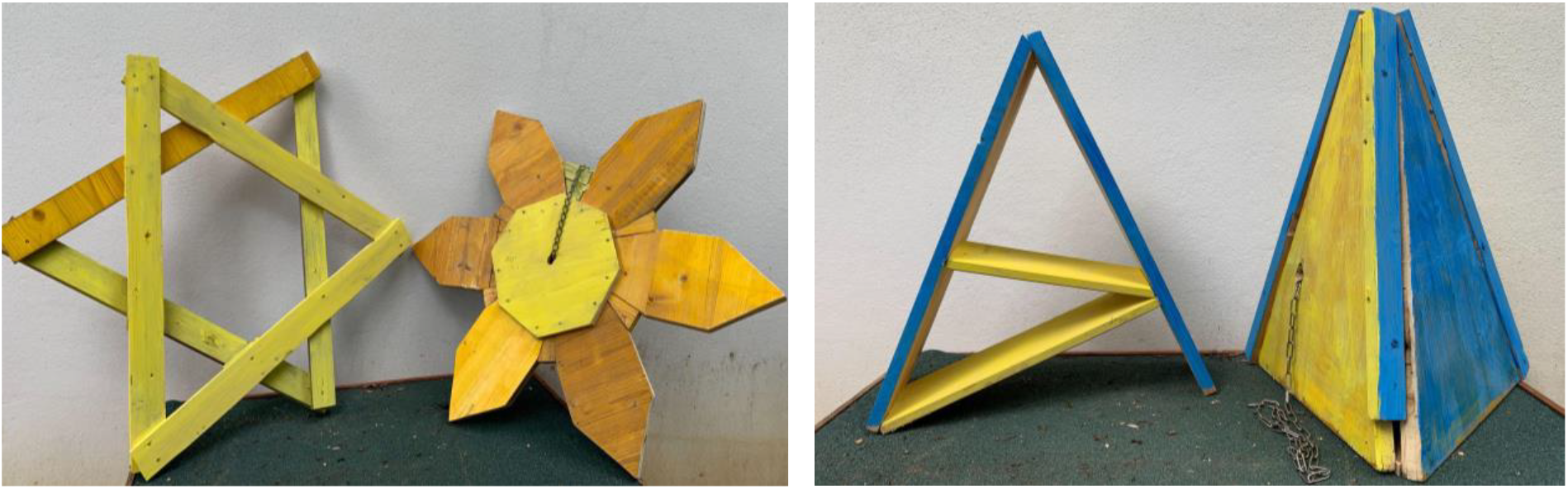
Example of object pairs used in the experiment. For the pictures of all objects (along with their descriptions, see Figure S1 in the supplementary materials.

We built 4 object pairs in total (see Figure S1 in the supplementary materials for the images of all the objects), and each animal was exposed to 3 of them during the course of the experiment. We counterbalanced the number of times each pair of objects would be used per population, as well as the order in which it would be presented (1^st^, 2^nd^ or 3^rd^).

For more details on what objects were used for what subjects, see Table S3.

### Cameras

The cameras used for filming the first reactions and the test sessions were Sony HDR-CX cameras.

We used camera traps for all the interactions with the object during the exposure phase (Browning trail cameras with night vision (1920×1080 resolution with Audio). We set the program IR smart vision, which allowed us to get videos at each detected movement regardless of the time of the day. We programmed them to detect movement up to 80 feet and to record for a minimum of 30 seconds.

### Experimental procedure

#### Exposure phase

##### Wolves and pack dogs

We exposed each pack to the respective object in their home enclosures. Each pack was exposed to one object for one week, with the object therefore becoming “familiar” for that pack in the test phase —in which the other object from the same pair (the “new” object) was presented as well.

We placed the object in the enclosure while the animals were away and placed it near one of their usual paths so that it would be more likely that the subjects would encounter it. The object was attached with a chain to a tree so that it could not be removed by the animals. The tree was chosen so that it was 10 meters apart from a second tree where the “new” object would be attached during the test phase. However, the two selected trees were not equidistant from the animals’ entrance due to the landscape differences between enclosures (see details table S1 in the Supplementary Materials; this was considered for the statistical analyses).

We determined the body length of each species by calculating the average of the length from the muzzle to the rear of our subjects. We used this estimate to draw two perimeters on the ground around both trees marked by using a few stones and bricks to indicate the correct distance and to maximize visibility while disturbing the animal’s usual environment as little as possible. The first circle was drawn at a distance of one body length (1,30m for the wolves, 85 cm for the dogs) around the object. The second circle was drawn at a distance of two body lengths (2,60m for the wolves, 1,70m for the dogs).

Once the objects were affixed to the trees, the animals were let back into the enclosure, while we filmed their first reactions towards the object from the outside. Moreover, all interactions between animals and objects during the exposure phase were video recorded from the outside of the enclosure by camera traps mounted on tripods, to observe the habituation of the animals towards the objects throughout the whole phase.

At the end of this exposure phase, the object was removed, and the pack was released for one day in the empty enclosure. We did this because we previously observed that the animals eventually get habituated over time to the presence of objects in their home enclosure, which would overly highlight the novelty of the “new” object. After this day, the test took place in the same enclosure.

The animals were exposed to 3 different object pairs in this manner (followed by their respective test sessions), separated by an interim of 3-4 weeks.

##### Pet dogs

According to the dogs’ relationships with each other, we assigned the six pet dogs into two groups. As with the wolves and pack dogs, the object pairs differed per group and changed between sessions.

For the exposure phase, we put the group into an unused enclosure (similar to the home enclosures of the pack animals) for approximately 2 hours per day, for 5 days. Due to the availability of the owners, the exposure days were not always consecutive, and the test was not always performed directly on the next day after the last exposure session (see Table S4).

Moreover, due to the different protocol we used for the pet dogs, we did not use camera-traps with them, and thus they were not included in the statistical analyses for the exposure phase.

#### Test phase

The animals were tested in the same enclosures used for the habituation. The day after the removal of the object, the pack was moved out of their home enclosure and put into the shifting system^1^ so that they could not see the enclosure, and the respective object pair was placed inside while the animals were away. The familiar object was positioned at the exact same place as during the exposure phase to avoid any changes that may disturb the animals’ perception. Indeed, the same object in a different position or placed in a different manner could be perceived by the animals as a different object (as noticed in previous experiments). The new object was attached by a chain to the second tree, located approximately 10 meters from the first one (See Figure 2). Once the set-up was ready, the subjects were separated from other pack members and tested individually (except NU in session 1 and 2 due to issues separating the animals and TT in session 1 due to experimental error; see Table S1 for more details).

**Figure 2:**
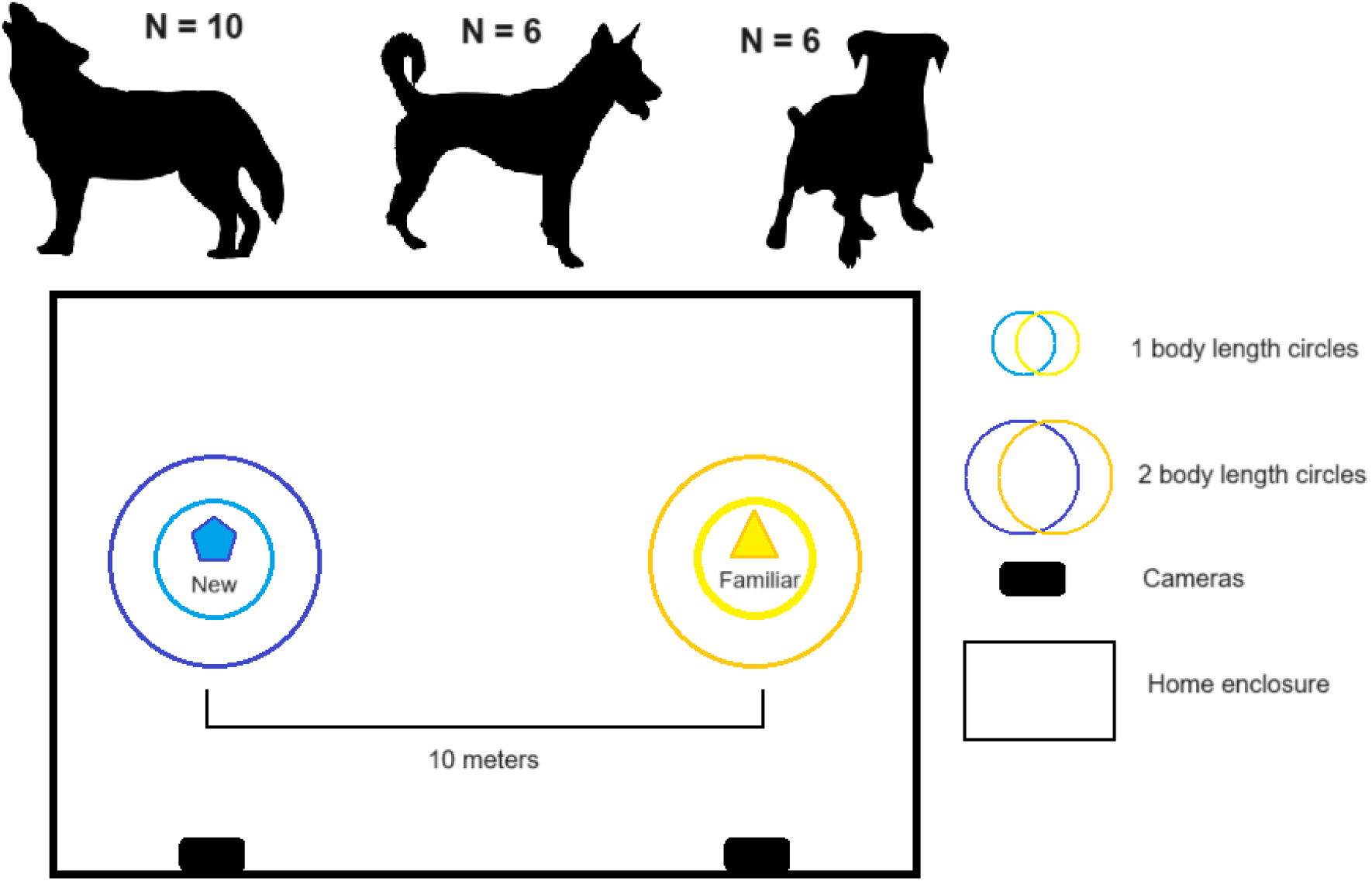
Experimental set-up for the test sessions. The three populations used in this study were, from left to right, wolves (N=10), pack dogs (N=6) and pet dogs (N=6). Both objects were chained to trees in the animals’ home enclosure, separated by a distance of approximately 10 meters. The familiar object was chained to the same tree that had been used for the exposure phase. In order to facilitate the process of calculating the animals’ distance towards the object, two concentric circles made of small, loosely distributed stones and bricks were placed at a distance of 1 body length of the animal (averaged at 1,30m for wolves and 85 cm for wolves) and 2 body lengths. Each session was filmed by two cameras —each of them pointing towards one of the objects— for later analyses.

The test started when the subject was led back into the enclosure and ended when they stopped interacting with the object for 5 consecutive minutes. If the subject did not approach any object at all for 10 minutes, we stopped the session and counted it as “no approach” for both objects. If the subject did not stop interacting with the object at any point, the session had a maximum duration of 20 minutes (although this, in fact, never happened). When the test was over, the animal was taken back into the shifting system, and another member of the same pack was shifted in to be tested. Once all members of a pack were tested, we removed the objects before releasing the pack back into their home enclosure. The object pair was cleaned and repainted after each session to avoid olfactory effects (knowing that some animals urinated on them or licked them).

Two cameras were placed outside the fence of the enclosure facing each of the objects. The entry of the animal in the enclosure was considered as the starting point for the testing session (see Figure 2).

### Data analyses

#### Video analysis: Exposure phase

For every exposure phase video taken with the camera traps, we coded whether an individual entered within the 1 or 2 body length circles around the object). We defined and animal “being within a circle” as them having at least their two forelegs inside the circle. We only coded the closest zone towards the object an animal had gotten in each of the videos: if the animal got within a distance of 2 body lengths to the object but no further, we coded “2_body_lengths”, if they approached further into a distance of 1 body length, we coded “1_body_length”, and finally, if they went closer and interacted directly with the object, we coded “touch” (see Table S5 for more details).

We also measured the latency to the first time they got within a distance of 2 body lengths to the object. To get this measure, we kept track of the time of the day (in hours and minutes) at which the object was presented, and subtracted this from the time at which the first video with an approach was filmed (as recorded by the camera trap). This measurement was rounded to the closest minute (i.e., if the animals approached the object in less than a minute after they were let into the enclosure, the time to approach would be coded as 0 hours and 0 minutes).

Finally, using a stopwatch, we calculated the duration of the interactions with the object for each video.

#### Video analysis: Test phase

Latencies and durations of interaction with each object were coded from the videos using BORIS (Behavioural Observations Research Interactive Software) developed by Olivier Friard, according to the Ethogram presented in Table 1. We coded also the first object touched (new/familiar) and the first approached (new/familiar) for each subject.

**Table 1.** Ethogram used to code the test videos of the experiment, including the different behaviors, their codes, modifiers, and descriptions as well as the codes and descriptions of the modifiers.

| Behaviors | Event | Modifiers | Description |
| --- | --- | --- | --- |
| Approach | point event | Identity of the object | The subject comes within 2 body lengths of an object while orienting their head towards it |
| Interaction start | point event | Identity of the object | The subject touches, sniffs, bites, or otherwise directly interacts with the object. |
| Interaction | duration | Identity of the object | The subject is interacting with an object as described above. Has to be preceded by “Interaction start” and bookended by “Interaction end”. |
| Interaction end | point event | Identity of the object | The subject stops interacting with the object (whether they moved away from it or not). |
| No approach | point event |  | The subject is in the object area but does not come within the circles. |

#### Statistical analyses

To evaluate if there was a significant difference between dogs’ and wolves’ behaviors, we performed several generalized linear mixed models (GLMMs, function glmer from the lme4 package) and linear mixed models (LMMs; lmer from lme4 (V.0.6.7, Bates et al., 2015) package) using R 4.0.2 (R Core Team, 2013). The predictor variable (fixed effect) was the group (wolf, pack dog, pet dog). In order to avoid making any direct comparisons between wolves and pet dogs (as their different upbringings and living conditions may have too big an influence on their behavior) and to avoid incurring in multiple testing, we chose pack dogs as the “intercept” in all of our models, and we did not run any analyses to compare wolves and pet dogs. The control variables (fixed effects) were the session (1, 2, 3), which object was closer to the entrance (new or familiar) and the distance from the entrance to the new object (in order to account for the different amount of time the animals would need to reach the object in each enclosure). The subject’s identity and the identity of each object were used as random effects in the different models. The degrees of freedom for the linear mixed models were calculated by using Satterthwaite’s method through the lmerTest (V.0.8.2, Kuznetsova et al., 2017) package.

Using the exposure phase data, we ran two LMMs: one for the time it took them to get within 2 body lengths of it and one for the duration of interaction with the object (in each video in which the subject was observed interacting with the object).

Due to technical difficulties with the camera-traps (battery life, storage space of the SD cards, malfunctioning due to extreme weather conditions, etc.), we did not always have a whole week (168 hours: 24 hours in 7 days) of camera trap functionality per pack and session (a full report on the estimated time the camera trap was functional per pack and session can be found on Table S6). To account for this, we used the number of hours the camera was recording as an offset in the model relating to the duration of interaction with the object. We did not include this measure for the model regarding the time the animals needed to reach within two body lengths of the object since these malfunctions occurred after the animals approached the object for the first time in all cases.

For the test phase, we ran GLMMs for each of the response variables using the test data: a model for the first object approached (omitting the instances in which no object was approached) and another one for the first object touched (omitting the instances in which no object was touched). We also ran a LMM for the durations of interaction with the familiar and the new object and a last one for the latency to approach the new object (omitting the instances in which the new object was not approached). For the duration of interaction, we added 1 to the logarithm of the initial duration because we had values equal to 0 in our data set (i.e., when the animals did not interact with one of the objects).

Random slopes were added whenever appropriate, and the model assumptions were met for each model. The model stability was analyzed through a custom function devised by Roger Mundry and modified by Remco Folkertsma, and the collinearity by using the “vif” function of the “car” (Fox and Weisberg, 2018) package. We compared all models with a null model lacking the predictor variables of the full model (i.e., population) but otherwise containing the same random effects and control variables. In all models, we z-transformed all continuous predictors (session) to obtain more easily interpretable model estimates and prevent model convergence issues. Then, we used anova function to compare the full-null models with the Chi-squared test. All models, including complete random-effects structure, are reported in the Table S7 in the Supplementary materials.

A second observer coded 20% of the videos and there was an agreement higher than 0.8 on all coded behaviors. The inter-observer reliability analyses showed that the duration of interaction (for both objects) had an intra-class correlation (ICC(_1_)) of 0.97 (F(_31,32_) = 74.2, p-value < 0.001), the latency to reach the new object had ICC(_1_) of 1, (F(_15,16_) = 13772, p< 0.001). Regarding the exposure phase data, the coding for whether or not an animal had interacted with the object had a Cohen’s Kappa (κ) equal to 0.89 (z = 42.1, p < 0.001), and the duration of interaction with the object had a ICC(_1_) of 0.98 (F(_119,120_) = 80, p < 0.001).

### Ethics

All dogs and wolves live in enclosures with their packs, have continuous access to water and are fed according to their individual needs. All procedures were approved by the “Ethics and Animal Welfare Committee at the University of Veterinary Medicine Vienna” with the ETK number ETK-079/05/2021.

## Results

### Exposure phase

All pack dogs and wolves approached to within 2 body lengths in all three exposure sessions. On average, wolves got within 2 body lengths of the object 0.162±0.105 hours after they were released into their enclosure, and pack dogs did so after 0.115±0.052 hours. Latency to the first approach within two body lengths was not significantly different between wolves and dogs (full-null comparison: χ^2^ = 0.008; p = 0.931; see Table S8 for the summary of the individual and population-wide values).

Most first approaches within two body lengths of the object (0,792%) led to an immediate interaction with the object, exception for 3 wolves who did not touch the object directly after approaching it in two sessions (*Una, Wamblee*, and *Kenai*),. Additionally, 2 wolves (*Nanuk and Yukon*) and 2 pack dogs (*Zuri* and *Enzi*) refrained from touching the object in only one session. Based on this observation, we decided not to conduct further analyses on the first approach within one body length or the first time the object was touched, as these measurements largely overlapped with the first approach within two body lengths in most cases.

Wolves interacted with the object for an average of 10.40 ± 0.69 seconds, while pack dogs did so for an average of 5.76 ± 0.56 seconds: This difference was significant (full-null comparison: χ^2^ = 6.37; p = 0.01); with the wolves interacting for longer with the object than the pack dogs (estimate ± SE = 0.49 ± 0.18, t(_27.09_) = 2.71, p-value = 0.01; see Table S9 for more details).

### Test phase

All pack dogs always interacted with at least one of the objects in every session, whereas pet dogs and wolves did not always do so. More specifically, 4 pet dogs (*Pepeo, Freya, Hakima*, and *Zazu*) failed to approach either object at least once and 3 wolves failed to approach either object at least once (*Nanuk, Una*, and Wamblee). When taking into account all the test sessions, wolves failed to approach either object in 13.3 ± 10.1% of the sessions and to touch them in 23.3 ± 13.6% of the sessions. Similarly, pet dogs failed to approach either object in 27.8 ± 18.8% of the sessions and to touch them in 33.3 ± 18.8% of them (Figure 3). However, since all pack dogs approached at least one of the objects in all sessions, we could not run a statistical model that tested the likelihood to approach (as one of the three levels of the predictor variable had no variation).

**Figure 3.**
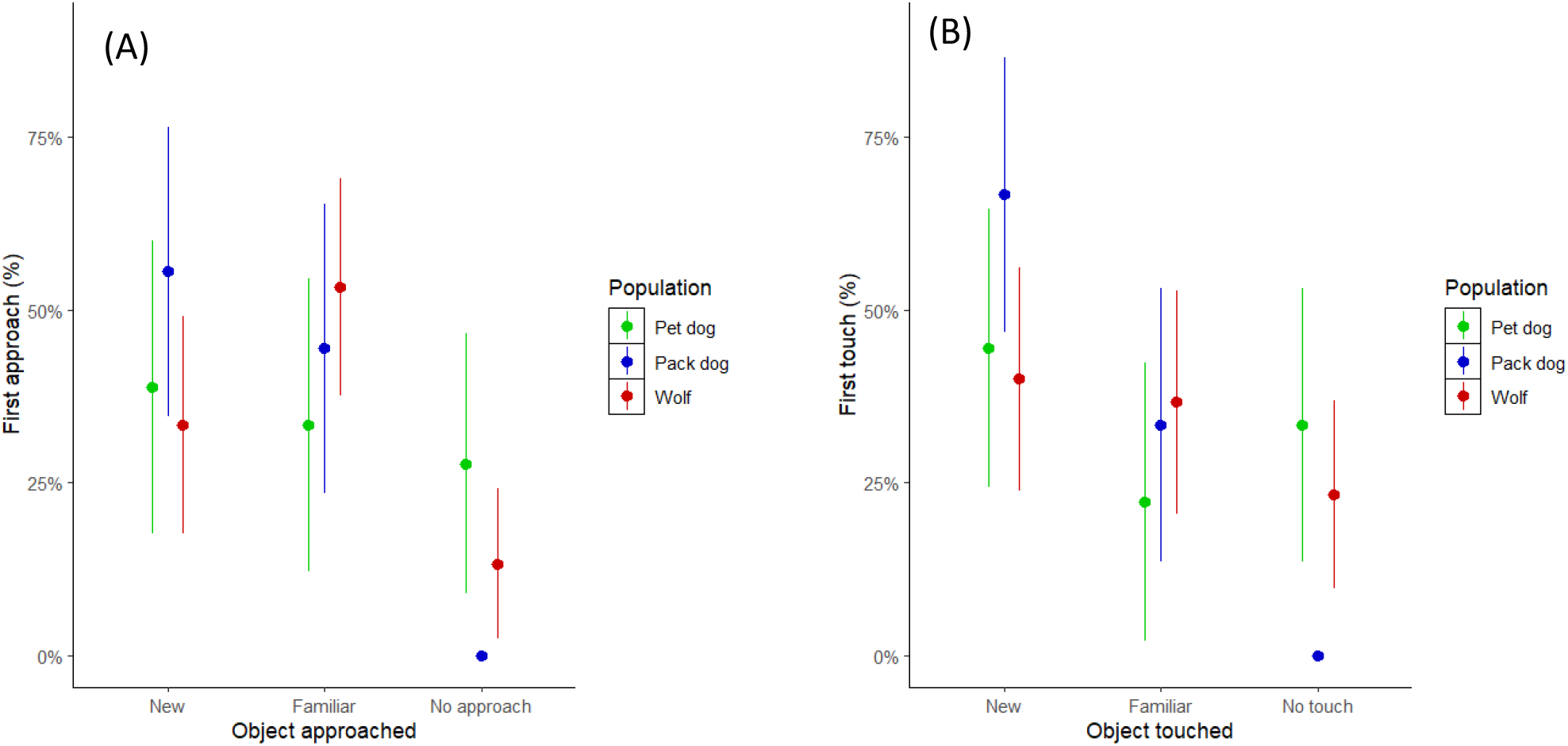
Percentage of sessions in which an animal chose to approach or touch the familiar or new object during the test. (A) Percentage of animals that approached the familiar or the new object first. (B) Percentage of animals that touched the familiar or the new object first. *From left to right in both plots: pet dogs (green), pack dogs (blue), and wolves (red)*.

Considering only sessions in which a subject approached at least one of the objects, we found that the wolves approached the new object first in 36.5% of the sessions (binomial test CI: 0.202 – 0.594, p = 0.3269), the packs dogs in 55.6% (binomial test CI: 0.308 – 0.785, p = 0.815), and the pet dogs in 53.9% (binomial test CI: 0.251 – 0.808, p = 1). This difference between populations was not significant (full-null comparison: χ^2^ = 1.05; p = 0.59).

In the sessions in which the animals touched at least one object, dogs’ preference was not significantly different from chance (pack dogs: 66.7% of the sessions, binomial test CI: 0.410 – 0.867, p = 0.238; pet dogs: 66.7% of the sessions, binomial test CI: 0-348 – 0.901, p = 0.388). Wolves did not present a preference towards interacting first with the new object either (52.2%, binomial test CI: 0.306 – 0.732, p = 1). Overall, we did not find any differences in the first object touched between wolves and pack dogs, nor between pack dogs and pet dogs (full-null comparison: χ^2^ = 4.61; p = 0.33). Thus, taken together, no population effect emerged in the preference towards a “new” vs. a “familiar” object.

Animals that approached one of the objects almost always interacted with it as well, with 93.0% of the sessions (53 out of 57) having a “first approach” followed by a “first touch”. Out of the instances in which both an approach and an interaction took place, there were only a few cases in which the first object approached was not the same as the first object touched (7 out of 70 —13.2%—, all of them being first approaches towards the familiar object, but first 392 interactions with the new one).

On average, latency to interact with the new object was 77.50 ± 29.80 seconds for pet dogs, 23.20 ± 8.02 seconds for pack dogs, and 33.50 ± 8.50 seconds for wolves. No significant differences between the groups emerged (full-null comparison: χ^2^ = 2.54; p = 0.28, see Figure 4).

**Figure 4.**
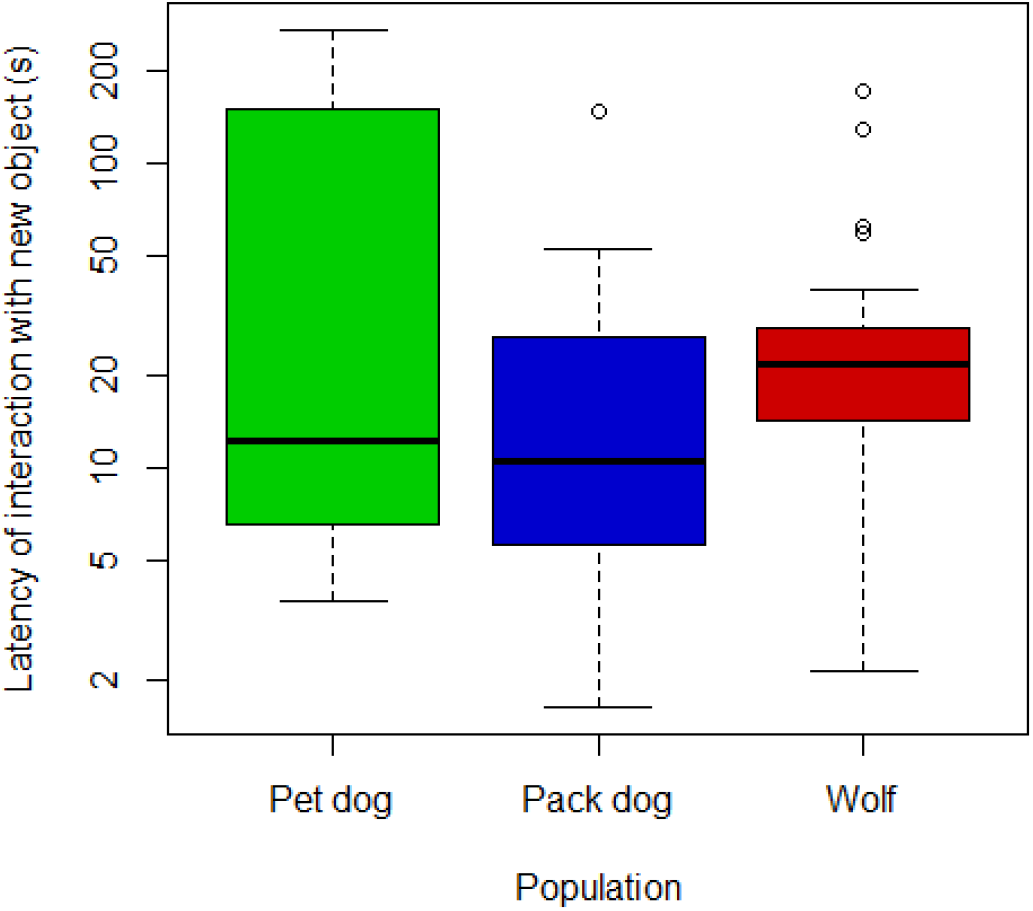
Latency to interact with the new object within the session in which the new object was interacted with, as a function of population. Latency values were log-transformed.

Duration of interaction with the familiar object in the test sessions was, on average, 1.79 ± 0.82 seconds for pet dogs, 4.66 ± 1.28 seconds for pack dogs, and 3.31 ± 0.97 seconds for wolves. In the case of the new object, pet dogs interacted with it an average of 3.74 ± 1.08 seconds, pack dogs an average of 10.70 ± 1.81 seconds, and wolves an average of 12.00 ± 2.80 seconds (see Figure 5 and Table S10). The full model for the duration of interaction with both objects was significantly different from the null (full-null comparison: χ^2^ = 25.40; p < 0.001). Since the interaction between the object (familiar/new) and population (i.e., whether each group of subjects interacted for longer with one of the objects over the other) was not significant (pack dogs vs. pet dogs: estimate ± SE = -0.03 ± 0.48, t(_54.00_) = -0.07, p = 0.94; pack dogs vs. wolves: estimate ± SE = 0.25 ± 0.43, t(_52.48_) = 0.59, p = 0.56), we ran a reduced version of the model that excluded said interaction. This model was significantly different from the null model as well (reduced-null comparison: χ^2^ = 24.91; p < 0.001). We found that the animals of all populations interacted significantly more with the new item than with the familiar one (estimate ± SE = 0.99 ± 0.17, t(_27.40_) = 5.85, p < 0.001). However, there were no significant differences in duration of interaction with either object between pack dogs and wolves (estimate ± SE = 0.29 ± 0.23, t(_10.71_)= 1.30, p = 0.22).

**Figure 5.**
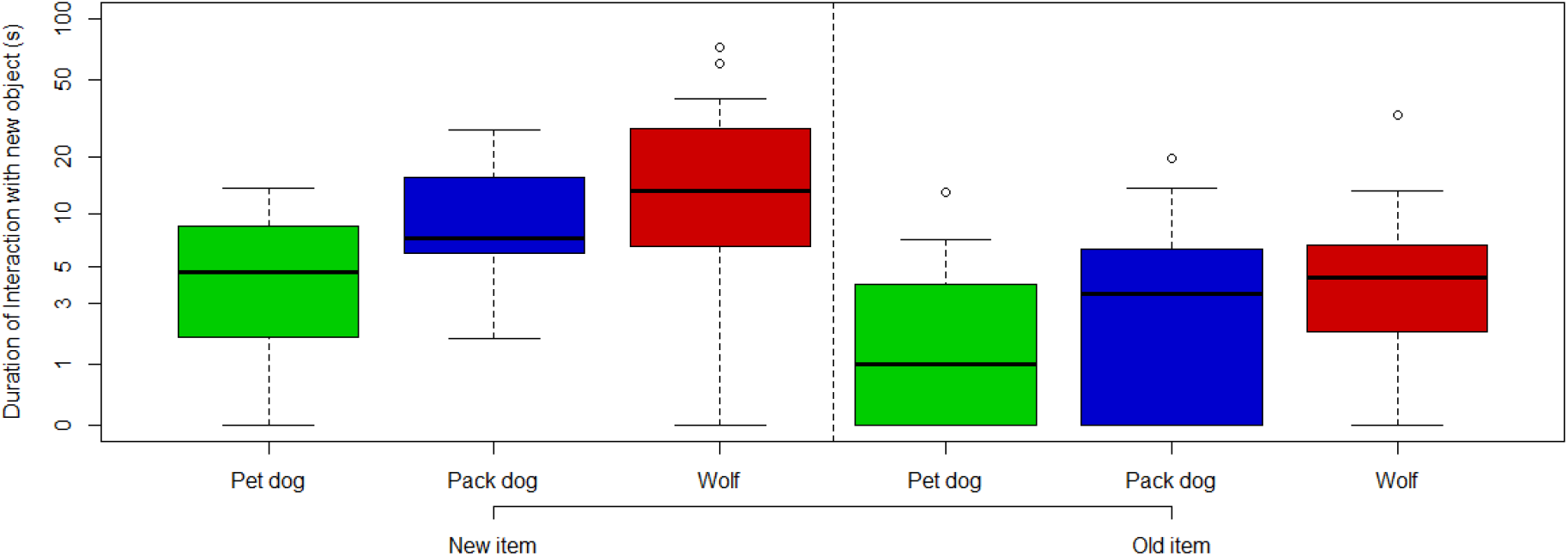
Graph comparing the duration of the interactions between familiar and new items per populations. Durations were added a value of 1 and later log-transformed (in order to avoid issues of trying to log-transform values of 0 whenever there was no interaction with one of the objects).

## Discussion

In the current study we aimed to test the effects of domestication on neophilia. To do so, we employed two different methods: first, the presentation of a novel object in the animals’ enclosure (exposure phase) and second, presenting this now ‘familiar’ object again together with another, novel one to investigate the animals’ potential preferences towards one of them (test phase). Our results showed a longer interaction time with the new object over the familiar one, but no differences between the populations in this measure.

In the exposure phase, wolves needed approximately the same time as pack dogs to approach the object. Wolves did, however, interact more with the familiar object during the exposure phase than the pack dogs. This result is partly in line with Moretti et al., (2015), who found that wolves took longer to approach the novel object but then, similarly to results in this study interacted longer with the object than dogs.

In the test phase however, we did not find any differences between pack dogs and wolves nor between pack dogs and pet dogs in the preference towards the new object and latency to approach it. It is important to note, however, that these analyses only took into account the sessions in which the animals approached one of the objects; something that was always the case for pack dog sessions, but not so for the pet dogs and the wolves (which failed to approach either object in around 30% of the trials; figure 3).

The results of the test phase seem to contrast with to those presented by Moretti et al. (2015) both in terms of latency to approach the novel object and the duration of interaction with it. Further, in the study by Moretti and colleagues it was dogs and not wolves that failed to approach the object in some of the trials, Nevertheless, it is important to take into consideration that the part of our experiment that is comparable with the one by Moretti and colleagues’ is the aforementioned exposure phase and, as such, such discrepancies may have come as a result of the differences in protocol (most notably, the fact that two objects were presented at once).

All three populations interacted for a longer amount of time with the novel item than with the familiar one. This could be due to reduced interest in the familiar item or to heightened interest in the new one. After one week of exposure to the familiar object, the animals would have ample opportunity to determine that interacting with it would provide no benefit (e.g., access to food). Conversely, the novelty of the new object could have driven them to interact with it for longer, in order to gather information about it and any opportunities that it may have been linked to (which could be considered a neophilic response). These two potential explanations are, of course, not mutually exclusive. The f unction of neophilia is to drive animals to gather information on the environment that they can exploit it later on (Griffin & Guez, 2014; Mettke-Hofmann et al., 2002).

Contrary to Kaulfuß and Mills’ (2008) hypothesis that neophilia in dogs is an adaptation for exploiting human environments, our findings showed no significant differences in neophilia between wolves and dogs. Both species appeared to favor the novel object over the familiar one, based on the time spent interacting with it. However, it is possible that the ancient dogs’ approach to human settlements was driven not by an increase in neophilia, but by a reduction in neophobia. That is to say, it is possible that ancient dogs were not more driven to explore novel features of the environment, but rather had a reduced fear response.

It is important to keep in mind, however, that life in captivity has been reported as a factor that would reduce animals’ neophobia and increase their neophilia (Benson-Aram et al., 2013; Damerius et al., 2017; Lazzaroni et al., 2019, Souganidis et al., 2024). In fact, our results fall in line with those described by Souganidis et al. (2024), in which captive individuals from several species of primates had a preference towards novel objects over familiar ones, regardless of the differences in feeding ecology between the species. Although we did test pet dogs in our study as a means to analyze the influence of experience in neophilia, all three of our populations lived in a human-shaped environment in which abundant food was provided and there was no risk of predation. Thus, it is possible that the living conditions of the pack-living wolves and dogs may have been enough to drive them towards neophilic behaviors. That being the case, using this paradigm the wild-living counterparts of our subjects —wild wolves and free-ranging dogs— would provide valuable insight into not only their potential differences in neophilia, but also on the effect of captivity in such measurements.

Measurements of neophilia and neophobia have been deemed as difficult to tease apart in the literature (Greenberg & Mettke-Hofmann, 2001). Although studies measuring neophobia usually rely on the presence of an item the animal may have an inherent drive to approach (usually food), measurements of neophilia do not have any particular constraints, as long as such external motivators are not involved. In the current study we did find that animals did interact more with the new item which could be considered a measurement of neophilia. However, it is ultimately not possible to determine the role other factors (such as neophobia) may have played in some of our measurements (e.g., the latency towards approaching the novel object). Thus, future studies should test neophilia (potentially through a paradigm similar to this one) as part of a test battery that includes other known factors that may have influenced our results (such as neophobia and general activity levels of each animal) in order to get a more precise measurement.

Aside for neophobia, other confounding factors can influence interspecific comparisons. Presenting the animals with several sets of objects and using the animal’s preference towards either the novel object or the familiar one as a measure of neophilia has the potential to mitigate the effect of some confounding factors. Some features of an item (size, shape, color) may be preferred or avoided by a particular individual or species, so habituating the animals to an item that has similar properties to the novel item has the potential to act as a “baseline”, as any preference in approaching and interacting with the novel object would only be explained by neophilia.

In conclusion, although we did not find any significant differences in neophilia between adult wolves and dogs, our paradigm could be used to gain further insights in wolves and dogs’ responses to novelty. Future studies should place a greater focus in trying to tease apart the individual effects of neophilia and neophobia, potentially through the use of neophobia tests as a baseline. Furthermore, testing wild-living populations should also provide valuable information on the ways the different species’ feeding ecology may drive differences in neophilia and neophobia, as well as the effect of life in captivity on such measurements.

## Supporting information

Dataset (exposure phase)

Dataset (test phase)

## Author contributions

FR and SMP conceptualized the experiment; LGDR and DRB developed the methodology and crafted the objects; LGDR and DRB performed the data collection; DRB and LGDR curated and analyzed the data; DRB and LGDR wrote the first draft of the paper; FR, ST and SMP supervised the entirety of the experiment; all authors edited and reviewed the manuscript.

## Acknowledgements

We thank Kathryn Roddie and the trainers at the Wolf Science Center Core facility for contributing to the data collection process.

## Funding statement

This research was funded by the Austrian Science Fund (FWF) DOI: 10.55776/P33928. DRB was funded by a Marietta Blau Grant from the Austrian Agency for International Cooperation in Education, Science and Research (MMC-2023-07030). For open access purposes, the authors have applied a CC BY public copyright license to any author-accepted manuscript version arising from this submission.

## Footnotes

1 All enclosures are surrounded by a fenced corridor where the animals can be placed in order to perform maintenance activities or move the animals to adjacent enclosures.

## Notes

### Competing Interest Statement

The authors have declared no competing interest.

## References

Bates, D., Mächler, M., Bolker, B., & Walker, S. (2015). Fitting Linear Mixed-Effects Models Using lme4. Journal of Statistical Software, 67(1), 1–48.

Benson-Amram, S., & Holekamp, K. E. (2012). Innovative problem solving by wild spotted hyenas. Proceedings of the Royal Society B: Biological Sciences, 279(1744), 4087–4095.

Benson-Amram, S., Weldele, M. L., & Holekamp, K. E. (2013). A comparison of innovative problem-solving abilities between wild and captive spotted hyaenas, Crocuta crocuta. Animal Behaviour, 85(2), 349–356.

Biondi, L. M., Bó, M. S., & Vassallo, A. I. (2010). Inter-individual and age differences in exploration, neophobia and problem-solving ability in a Neotropical raptor (Milvago chimango). Animal cognition, 13, 701–710.

Butler, J. R., Brown, W. Y., & Du Toit, J. T. (2018). Anthropogenic food subsidy to a commensal carnivore: the value and supply of human faeces in the diet of free-ranging dogs. Animals, 8(5), 67.

Brooks, R. (1990). Survey of the dog population of Zimbabwe and its level of rabies vaccination. The Veterinary Record, 127(24), 592–596.

Cooper, A., Turney, C., Hughen, K. A., Brook, B. W., McDonald, H. G., & Bradshaw, C. J. (2015). Abrupt warming events drove Late Pleistocene Holarctic megafaunal turnover. Science, 349(6248), 602–606.

Crane, A. L., & Ferrari, M. C. (2017). Patterns of predator neophobia: a meta-analytic review. Proceedings of the Royal Society B: Biological Sciences, 284(1861), 20170583.

Damerius, L. A., Forss, S. I. F., Kosonen, Z. K., Willems, E. P., Burkart, J. M., Call, J., Galdikas, B. M. F., Liebal, K., Haun, D. B. M., & van Schaik, C. P. (2017). Orientation toward humans predicts cognitive performance in orangutans. Scientific Reports, 7(1), 40052.

Dufresnes, C., Miquel, C., Remollino, N., Biollaz, F., Salamin, N., Taberlet, P., & Fumagalli, L. (2018). Howling from the past: historical phylogeography and diversity losses in European grey wolves. Proceedings of the Royal Society B, 285(1884), 20181148.

Freedman, A. H., & Wayne, R. K. (2017). Deciphering the origin of dogs: From fossils to genomes. Annual Review of Animal Biosciences, 5(1), 281–307.

Fox, J., & Weisberg, S. (2018). An R companion to applied regression. Sage publications. Greenberg, Russell, and Claudia Mettke-Hofmann. “Ecological aspects of neophobia and neophilia in birds.” Current ornithology (2001): 119–178.

Greggor, A. L., Clayton, N. S., Fulford, A. J., & Thornton, A. (2016a). Street smart: faster approach towards litter in urban areas by highly neophobic corvids and less fearful birds. Animal Behaviour, 117, 123–133.

Greggor, A. L., Jolles, J. W., Thornton, A., & Clayton, N. S. (2016b). Seasonal changes in neophobia and its consistency in rooks: the effect of novelty type and dominance position. Animal behaviour, 121, 11–20.

Griffin, A. S., & Guez, D. (2014). Innovation and problem solving: a review of common mechanisms. Behavioural Processes, 109, 121–134.

Griffin, A. S., Tebbich, S., & Bugnyar, T. (2017). Animal cognition in a human-dominated world. Animal Cognition, 20, 1–6.

Heinrich, B. (1995). Neophilia and exploration in juvenile common ravens, Corvus corax. Animal Behaviour, 50(3), 695–704.

Kuznetsova, A., Brockhoff, P. B., & Christensen, R. H. B. (2017). lmerTest package: tests in linear mixed effects models. Journal of statistical software, 82(13).

Kaulfuß, P., & Mills, D. S. (2008). Neophilia in domestic dogs (Canis familiaris) and its implication for studies of dog cognition. Animal cognition, 11, 553–556.

Lazzaroni, M., Range, F., Bernasconi, L., Darc, L., Holtsch, M., Massimei, R., Rao, A., & Marshall-Pescini, S. (2019). The role of life experience in affecting persistence: A comparative study between free-ranging dogs, pet dogs and captive pack dogs. PLoS One, 14(4), e0214806.

Marshall-Pescini, S., Cafazzo, S., Virányi, Z., & Range, F. (2017a). Integrating social ecology in explanations of wolf–dog behavioral differences. Current Opinion in Behavioral Sciences, 16, 80–86.

Marshall-Pescini, S., Virányi, Z., Kubinyi, E., & Range, F. (2017b). Motivational factors underlying problem solving: Comparing wolf and dog puppies’ explorative and neophobic behaviors at 5, 6, and 8 weeks of age. Frontiers in Psychology, 8, 180.

Mech, L. D., Smith, D. W., & MacNulty, D. R. (2015). Wolves on the hunt: the behavior of wolves hunting wild prey. University of Chicago Press.

Mettke-Hofmann, C., Winkler, H., & Leisler, B. (2002). The significance of ecological factors for exploration and neophobia in parrots. Ethology, 108(3), 249–272.

Miranda, A. C., Schielzeth, H., Sonntag, T., & Partecke, J. (2013). Urbanization and its effects on personality traits: a result of microevolution or phenotypic plasticity?. Global change biology, 19(9), 2634–2644.

Moretti, L., Hentrup, M., Kotrschal, K., & Range, F. (2015). The influence of relationships on neophobia and exploration in wolves and dogs. Animal Behaviour, 107, 159–173.

Range, F., & Marshall-Pescini, S. (2022). Comparing wolves and dogs: current status and implications for human ‘self-domestication’. Trends in Cognitive Sciences, 26(4), 337–349.

Rao, A., Bernasconi, L., Lazzaroni, M., Marshall-Pescini, S., & Range, F. (2018). Differences in persistence between dogs and wolves in an unsolvable task in the absence of humans. PeerJ, 6, e5944.

R Core Team. “ R: A language and environment for statistical computing.” Foundation for Statistical Computing, Vienna, Austria (2013).

Sih, A., & Del Giudice, M. (2012). Linking behavioural syndromes and cognition: a behavioural ecology perspective. Philosophical Transactions of the Royal Society B: Biological Sciences, 367(1603), 2762–2772.

Singletary, M., & Lazarowski, L. (2021). Canine special senses: considerations in olfaction, vision, and audition. Veterinary Clinics: Small Animal Practice, 51(4), 839–858.

Souganidis, C., Llorente, M., Aureli, F., Call, J., & Amici, F. (2024). Variation in neophilia in seven primate species. Journal of Comparative Psychology.

Suzuki, K., Ikebuchi, M., Kagawa, H., Koike, T., & Okanoya, K. (2021). Effects of domestication on neophobia: A comparison between the domesticated Bengalese finch (Lonchura striata var. domestica) and its wild ancestor, the white-rumped munia (Lonchura striata). Behavioural Processes, 193, 104502.

Tebbich, S., Griffin, A. S., Peschl, M. F., & Sterelny, K. (2016). From mechanisms to function: an integrated framework of animal innovation. Philosophical Transactions of the Royal Society B: Biological Sciences, 371(1690), 20150195.

