## Supplementary material (tables and figures) for "Neophilia in wolves and dogs"

### Supplementary materials

Within wolves, we observed that three individuals with Eurasian ancestry (*Maikan, Tekoa, and Taima*) seemingly took more time than the rest of the population to approach the object in the exposure phase, and only one of them (*Tekoa*) touched it once. During the test sessions, none of them approached nor touched any of the items. Due to this behavior, we determined they did not complete the requisites to participate in the test (i.e., being habituated to the familiar object) and thus, we excluded them from the analyses.

(A)

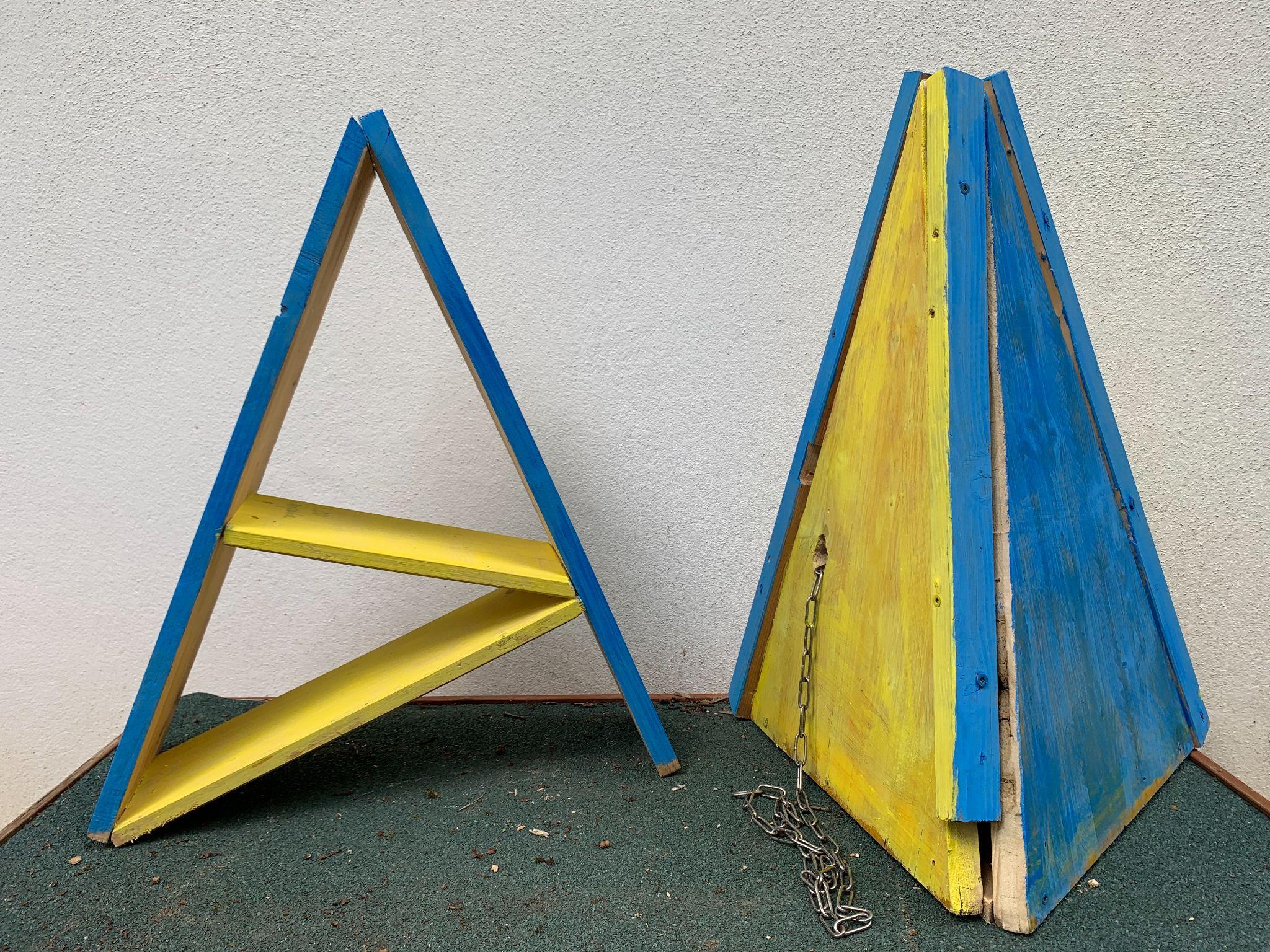

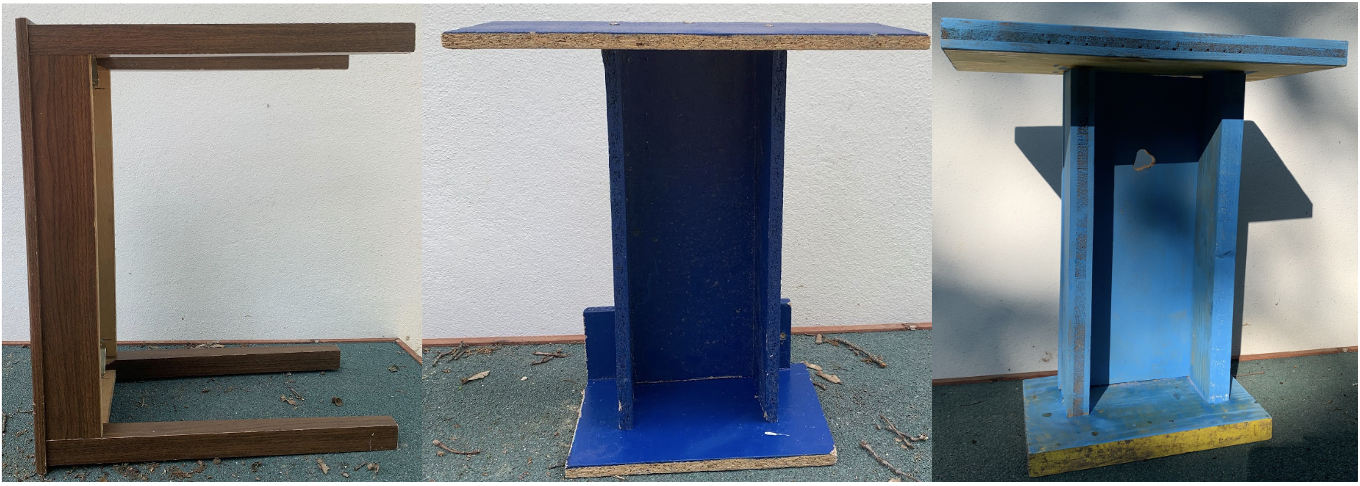
(B)	(C)

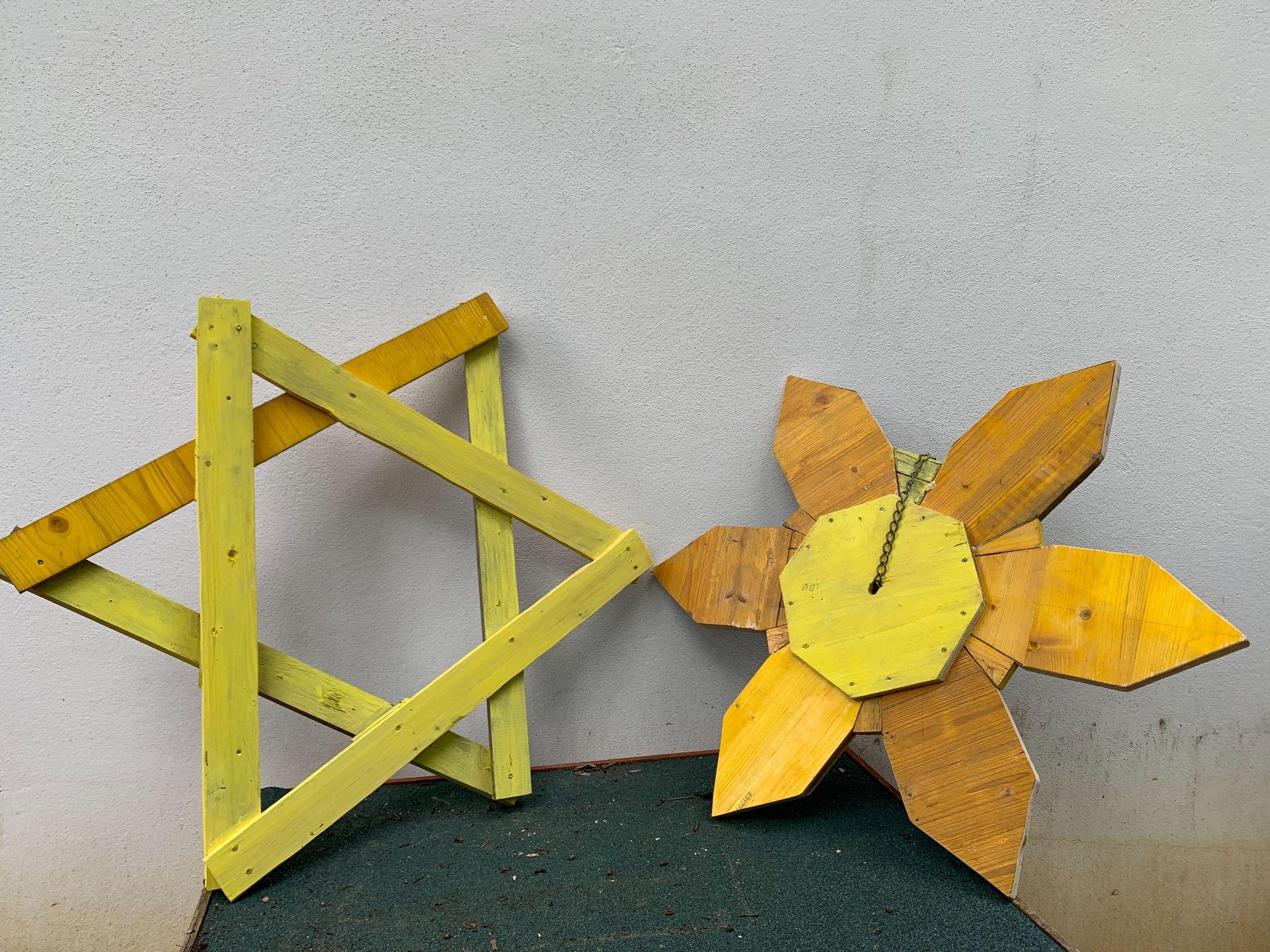

(D)

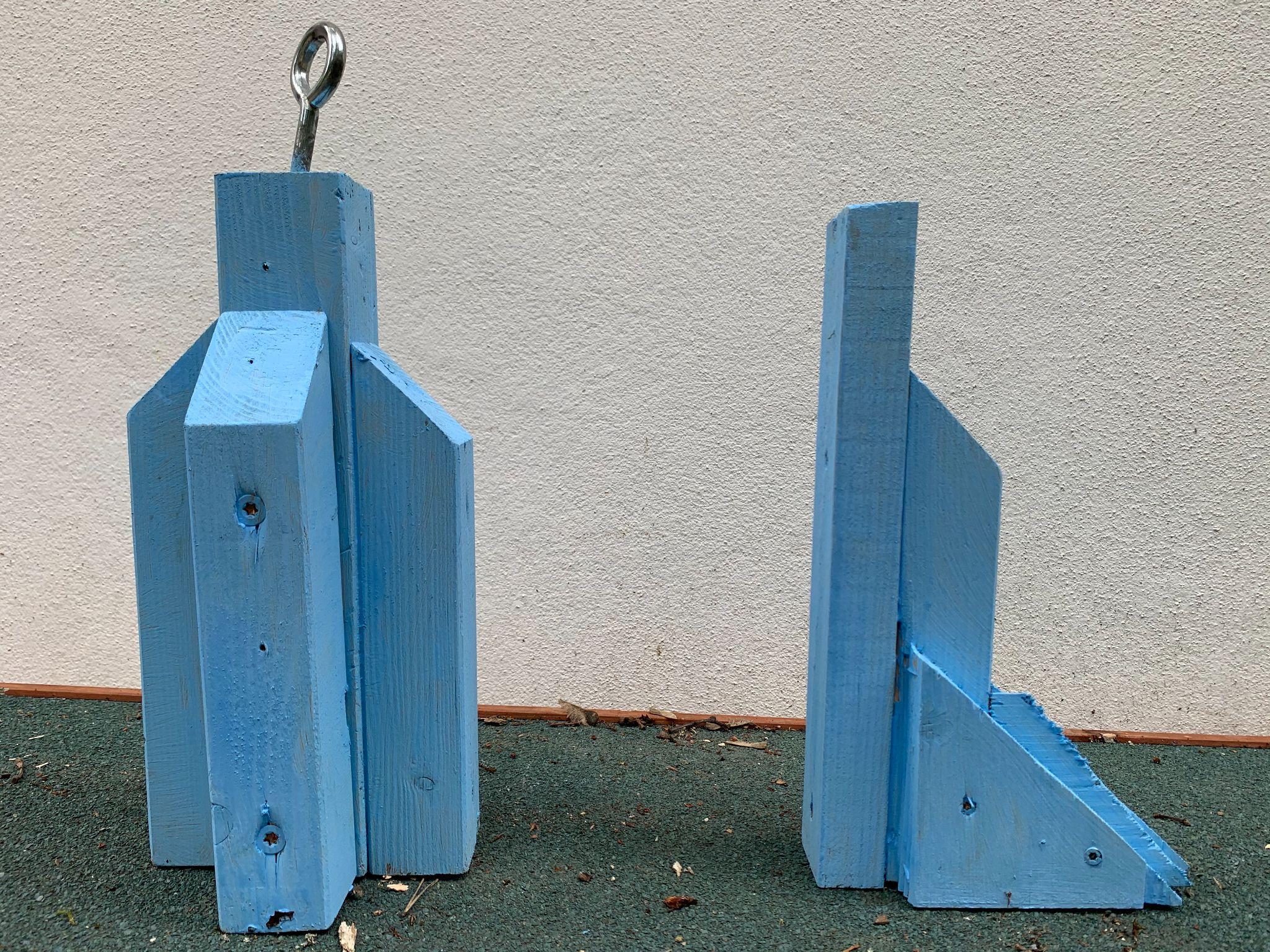

*Figure S1.* The different object pairs. The pair (A) was composed of a brown table and a dark blue T-shape and was approximately 50 cm high. A similar T-shape was built during the experiment because the first one was damaged by the wolves during a habituation phase (after which this pair was re-named as “E”). The pair (B) was composed of a star and a flower both painted in yellow and was 1 m high. The pair (C) was composed of a pyramid and a capital A both painted in yellow and blue and was 65 cm high. Finally, the pair (D) was composed of a rocket and a pistol both painted in light blue and was 30 cm high.

*Table S1.* Description of the enclosures and distribution of objects and trees during the experiment.

| Enclosure | Size (m^2^) | Closer object from entrance | Distance from entrance New | Distance from entrance Familiar |
| --- | --- | --- | --- | --- |
| 6 | 1923 | New | 13,1 | 20,8 |
| 9b | 10364 | Familiar | 54,1 | 51,7 |
| 9c | 1258 | Wolves: New Pet dogs: Familiar | Wolves: 7,3  Pet dogs: 13,7 | Wolves: 9,3  Pet dogs: 10,1 |
| 10a back | 2018 | Familiar | 17,9 | 4,5 |
| 10b front | 2014 | New | 14,4 | 11 |
| 10b back | 2012 | Familiar | 18,2 | 9,2 |
| 21 | 4043 | New | 12,6 | 22,6 |
| 22 | 4005 | Familiar | 25,4 | 15,4 |
| 23 | 2039 | Familiar | 13,1 | 8,4 |
| 24 | 2086 | Familiar | 26,5 | 14,5 |
| OTE | 400 | New | 10,6 | 12,6 |

*Table S2*. Summary table of the individuals tested, including their pack, age at the beginning of the experiment, origin, species, owners, relatives, and sex. Subjects greyed out were not considered for data analyses.

| Subject | Pack / Group | Species / Population | Subspecies / Breed | Owner | Origin | Relatives | Age (months) | Sex |
| --- | --- | --- | --- | --- | --- | --- | --- | --- |
| Taima | TT | Wolf | Timber + Eurasian wolf | WSC | Moscow | Tekoa, Maikan | 62 | Female |
| Tekoa |  |  | Timber + Eurasian wolf |  | Moscow | Taima, Maikan | 62 | Male |
| Nanuk | NU |  | Timber wolf |  | Triple D Game Farm, Montana, USA |  | 147 | Male |
| Una |  |  | Timber wolf |  | Minnesota Wildlife Connection, USA | Chitto | 111 | Male |
| Geronimo | GYW |  | Timber wolf |  | Triple D Game Farm, Montana, USA | Yukon | 146 | Female |
| Yukon |  |  | Timber wolf |  | Triple D Game Farm, Montana, USA | Geronimo | 146 | Male |
| Wamblee |  |  | Timber wolf |  | Haliburton Forest & Wildlife Reserve, Ontario, Canada | Etu | 111 | Female |
| Tala | TC |  | Timber wolf |  | Minnesota Wildlife Connection, USA | Amarok | 111 | Male |
| Chitto |  |  | Timber wolf |  | Minnesota Wildlife Connection, USA | Una | 111 | Female |
| Maikan | ME |  | Timber + Eurasian wolf |  | Moscow | Tekoa, Taima | 62 | Male |
| Etu |  |  | Timber wolf |  | Haliburton Forest & Wildlife Reserve, Ontario, Canada | Wamblee | 62 | Male |
| Amarok | AK |  | Timber wolf |  | Minnesota Wildlife Connection, USA | Tala | 111 | Male |
| Kenai |  |  | Hudson Bay wolf |  | Parc Safari Hemmingford, Quebec, Canada |  | 136 | Male |
| Hiari | IH | Pack dog | Mongrel |  | Wildpark Ernstbrunn, Austria | Imara | 88 | Male |
| Imara |  |  | Mongrel |  | Wildpark Ernstbrunn, Austria | Hiari | 88 | Female |
| Enzi | ZE |  | Mongrel |  | Wildpark Ernstbrunn, Austria | Layla, Panya, Zazu, Pepeo | 87 | Male |
| Zuri |  |  | Mongrel |  | Shelter Paks, Hungary |  | 121 | Female‡ |
| Layla | LP |  | Mongrel |  | Next to highway Györ, Hungary | Enzi, Panya, Zazu, Pepeo | 119 | Female |
| Panya |  |  | Mongrel |  | Wildpark Ernstbrunn, Austria | Layla, Enzi, Zazu, Pepeo | 87 | Female† |
| Asali | Group A | Pet dog | Mongrel | Owner 1 | Tierheim Szeged, Hungary |  | 130 | Male |
| Kilio |  |  | Mongrel |  | Shelter Paks, Hungary |  | 139 | Male* |
| Zazu |  |  | Mongrel | Owner 2 | Wildpark Ernstbrunn, Austria | Layla, Enzi, Panya, Pepeo | 87 | Male* |
| Hakima | Group B |  | Mongrel | Owner 4 | Shelter Paks, Hungary |  | 130 | Male* |
| Pepeo |  |  | Mongrel | Owner 3 | Wildpark Ernstbrunn, Austria | Layla, Enzi, Panya, Zazu | 87 | Male |
| Freya |  |  | Irish Terrier |  | Galahad’s Guardian Irish Terrier, Kirchberg, Switzerland |  | 74 | Female |

* Castrated.
† Went through false pregnancy on 11/10/2021.
‡ Spayed during the experiment (on 26/10/2021).

*Table S3*. Summary table of the usage of objects to the different animals, with their dates. Subjects greyed out were not considered for data analyses.

| Subject | Week item 1 | Pair 1 | Familiar Item 1 | New Item 1 | Week Item 2 | Pair 2 | Familiar item 2 | New Item 2 | Week Item 3 | Pair 3 | Old Item 3 | New Item 3 |
| --- | --- | --- | --- | --- | --- | --- | --- | --- | --- | --- | --- | --- |
| Tekoa | 02/08/2021 | A | T shape | Table | 27/09/2021 | B | Flower | Star | 22/11/2021 | C | Pyramid | Capital A |
| Taima | 02/08/2021 | A | T shape | Table | 27/09/2021 | B | Flower | Star | 22/11/2021 | C | Pyramid | Capital A |
| Nanuk | 02/08/2021 | C | Capital A | Pyramid | 13/09/2021 | D | Rocket | Pistol | 11/10/2021 | B | Star | Flower |
| Una | 02/08/2021 | C | Capital A | Pyramid | 13/09/2021 | D | Rocket | Pistol | 11/10/2021 | B | Star | Flower |
| Geronimo | 16/08/2021 | C | Capital A | Pyramid | 11/10/2021 | E | New T shape | Table | 08/11/2021 | B | Flower | Star |
| Yukon | 16/08/2021 | C | Capital A | Pyramid | 11/10/2021 | E | New T shape | Table | 08/11/2021 | B | Flower | Star |
| Wamblee | 16/08/2021 | C | Capital A | Pyramid | 11/10/2021 | E | New T shape | Table | 08/11/2021 | B | Flower | Star |
| Tala | 30/08/2021 | A | T shape | Table | 27/09/2021 | D | Pistol | Rocket | 08/11/2021 | C | Pyramid | Capital A |
| Chitto | 30/08/2021 | A | T shape | Table | 27/09/2021 | D | Pistol | Rocket | 08/11/2021 | C | Pyramid | Capital A |
| Maikan | 30/08/2021 | B | Flower | Star | 25/10/2021 | C | Capital A | Pyramid | 22/11/2021 | F | Table | New T shape |
| Etu | 30/08/2021 | B | Flower | Star | 25/10/2021 | C | Capital A | Pyramid | 22/11/2021 | E | Table | New T shape |
| Amarok | 13/09/2021 | B | Flower | Star | 25/10/2021 | E | New T shape | Table | 06/12/2021 | D | Pistol | Rocket |
| Kenai | 13/09/2021 | B | Flower | Star | 25/10/2021 | E | New T shape | Table | 06/12/2021 | D | Pistol | Rocket |
| Hiari | 02/08/2021 | B | Star | Flower | 13/09/2021 | C | Pyramid | Capital A | 22/11/2021 | D | Pistol | Rocket |
| Imara | 02/08/2021 | B | Star | Flower | 13/09/2021 | C | Pyramid | Capital A | 22/11/2021 | D | Pistol | Rocket |
| Enzi | 16/08/2021 | D | Rocket | Pistol | 27/09/2021 | E | Table | New T shape | 25/10/2021 | B | Star | Flower |
| Zuri | 16/08/2021 | D | Rocket | Pistol | 27/09/2021 | F | Table | New T shape | 25/10/2021 | B | Star | Flower |
| Layla | 30/08/2021 | D | Rocket | Pistol | 11/10/2021 | C | Pyramid | Capital A | 08/11/2021 | E | Table | New T shape |
| Panya | 30/08/2021 | D | Rocket | Pistol | 11/10/2021 | C | Pyramid | Capital A | 08/11/2021 | E | Table | New T shape |
| Asali | 16/08/2021 | A | Table | T shape | 11/10/2021 | D | Rocket | Pistol | 06/12/2021 | C | Capital A | Pyramid |
| Kilio | 16/08/2021 | A | Table | T shape | 11/10/2021 | D | Rocket | Pistol | 06/12/2021 | C | Capital A | Pyramid |
| Zazu | 16/08/2021 | A | Table | T shape | 11/10/2021 | D | Rocket | Pistol | 06/12/2021 | C | Capital A | Pyramid |
| Hakima | 16/08/2021 | A | Table | T shape | 01/11/2021 | D | Pistol | Rocket | 05/02/2022 | B | Star | Flower |
| Pepeo | 12/09/2021 | C | Capital A | Pyramid | 01/11/2021 | D | Pistol | Rocket | 05/02/2022 | B | Star | Flower |
| Freya | 12/09/2021 | C | Capital A | Pyramid | 01/11/2021 | D | Pistol | Rocket | 05/02/2022 | B | Star | Flower |

*Table S4*. Summary table of the exposure time pet dogs had per session.

| Exposure session | Subject group | Day 1 | Day 2 | Day 3 | Day 4 | Day 5 | Enclosure |
| --- | --- | --- | --- | --- | --- | --- | --- |
| 1 | Zazu, Kilio, Asali, Hakima^[1]^ | 1h40 | 2h30 | 2h15 | 1h15 | 1h45 | 24 |
|  | Pepeo, Freya | 2h50 | 1h50 | 2h10 |  |  | 9c |
| 2 | Zazu, Kilio, Asali | 1h10 | 2h15 | 2h30 | 2h15 | 2h00 | 10b_back |
|  | Pepeo, Freya, Hakima | 1h15 | 1h50 | 1h50 | 1h15 | 1h15 | 9c, then 6^[2]^ |
| 3 | Zazu, Kilio, Asali | 45min | 1h00 | 45min | 45min | 40min | OTE |
|  | Pepeo, Freya, Hakima | 50min | 45min |  |  | 50min |  |

*Table S5.* Code used to analyze the videos taken by camera-traps during the Exposure phase.

| Code | Behavior |
| --- | --- |
| 2_body_length | the animal enters the 2 body length circle |
| 1_body_length | the animal enters the 1 body length circle |
| nothing | nothing related to the experiment is on the video |
| touch | the animal sniffs, licks, grabs, pulls, uses his paw to touch the object |

*Table S6*. Amount of hours of videos we have recorded per pack and per session after we removed the time the videos were not available.

| Pack | Session | Amount of hours of videos |
| --- | --- | --- |
| TT | 1 | 168 |
|  | 2 | 168 |
|  | 3 | 89 |
| NU | 1 | 168 |
|  | 2 | 96 |
|  | 3 | 144 |
| GYW | 1 | 72 |
|  | 2 | 168 |
|  | 3 | 138 |
| ME | 1 | 168 |
|  | 2 | 72 |
|  | 3 | 156 |
| AK | 1 | 168 |
|  | 2 | 168 |
|  | 3 | 66 |
| LP | 1 | 168 |
|  | 2 | 168 |
|  | 3 | 168 |
| IH | 1 | 168 |
|  | 2 | 168 |
|  | 3 | 144 |
| ZE | 1 | 168 |
|  | 2 | 168 |
|  | 3 | 168 |

*Table S7.* The different models run during the statistical analysis of the study.

| **Test models** | |
| --- | --- |
| Function | Model structure |
| glmer | first_object_approached ~ population + z.session + closer_item + (1+z.session\|\|subject), family = "binomial" |
| glmer | first_object_touched ~ pop_full + z.session + closer_item + (1+z.session\|\|subject), family = "binomial" |
| lmer | log1p.duration_interaction ~ item * population + z.session + (1\|\|subject) + (1\|\|object_type), REML=F |
| lmer | log.latency_new ~ population + z.session + log.distance_new + (1+z.session\|\|subject), REML=F |
| **Exposure models** | |
| Function | Model structure |
| lmer | log.hours_to_habituate ~ population + z.session + (1+z.session\|\|subject) + (1+z.session+population.Wolf\|\|familiar_item), REML=F |
| lmer | log.duration_interaction ~ population * z.session + (1+z.session\|\|subject) + (1+z.session+population.Wolf\|\|familiar_item) + offset(log(hours_camera_trap_active)) , REML=F |

*Table S8*. Average amount of time (in hours) for every animal to approach within two body lengths to the familiar object.

| Subject | Mean latency to approach within two body lengths (hours) | Population | Mean per population |
| --- | --- | --- | --- |
| Amarok | 0.01±0.01 | Wolf | 0.16±0.05 |
| Chitto | 0.00±0.00 |  |  |
| Etu | 0.00±0.00 |  |  |
| Geronimo | 0.00±0.00 |  |  |
| Kenai | 0.00±0.00 |  |  |
| Nanuk | 0.00±0.00 |  |  |
| Tala | 0.02±0.02 |  |  |
| Una | 0.34±0.15 |  |  |
| Wamblee | 1.24±0.93 |  |  |
| Yukon | 0.00±0.00 |  |  |
| Enzi | 0.34±0.18 | Pack Dog | 0.12±0.05 |
| Hiari | 0.00±0.00 |  |  |
| Imara | 0.02±0.02 |  |  |
| Layla | 0.00±0.00 |  |  |
| Panya  Zuri | 0.00±0.00  0.34±0.18 |  |  |

*Table S9*. Mean duration of interaction (in seconds) with the object during the exposure phase per subject and per population.

| Subject | Mean duration of interaction | Population | Mean per population |
| --- | --- | --- | --- |
| Amarok | 7.62±1.85 | Wolf | 10.40±0.69 |
| Chitto | 12.50±3.26 |  |  |
| Etu | 17.60±2.57 |  |  |
| Geronimo | 7.64±1.52 |  |  |
| Kenai | 7.90±1.21 |  |  |
| Nanuk | 10.70±1.18 |  |  |
| Tala | 17.00±3.47 |  |  |
| Una | 5.76±1.14 |  |  |
| Wamblee | 8.67±2.38 |  |  |
| Yukon | 10.7±1.91 |  |  |
| Enzi | 5.59±1.26 | Pack Dog | 5.76±0.56 |
| Hiari | 5.35±1.07 |  |  |
| Imara | 6.10±1.10 |  |  |
| Layla | 4.30±1.08 |  |  |
| Panya  Zuri | 6.89±1.61  5.33±2.19 |  |  |

*Table S10*. Summary of the interaction durations of the different populations with the two objects in seconds.

| Population | Session | Mean duration interaction familiar object | Mean duration interaction familiar object per population | Mean duration interaction new object | Mean duration interaction new object per population |
| --- | --- | --- | --- | --- | --- |
| Pet Dogs | 1 | 2.50±1.24 | 1.79±0.82 | 5.39±1.07 | 3.74±1.08 |
|  | 2 | 0.38±0.38 |  | 0.00±0.00 |  |
|  | 3 | 2.50±2.20 |  | 5.83±2.57 |  |
| Pack Dogs | 1 | 6.59±1.88 | 4.66±1.28 | 9.31±2.09 | 10.70±1.81 |
|  | 2 | 4.04±3.23 |  | 14.50±4.17 |  |
|  | 3 | 3.33±1.28 |  | 8.29±2.64 |  |
| Wolves | 1 | 3.03±1.01 | 3.31±0.97 | 13.80±5.73 | 12.00±2.80 |
|  | 2 | 1.61±0.72 |  | 10.50±3.16 |  |
|  | 3 | 5.27±2.54 |  | 11.70±5.70 |  |

1. Hakima's first session was done with the first group due to the owner availability.
2. GYW pack had to be temporarily separated due intra-pack conflict; the enclosure where the pet dogs were habituated in had to be requisitioned for the wolves, thus we changed the enclosure for the pet dogs’ session.
